# Trade-off and flexibility in the dynamic regulation of the cullin-RING ubiquitin ligase repertoire

**DOI:** 10.1101/168898

**Authors:** Ronny Straube, Dietrich Flockerzi, Dieter A. Wolf

## Abstract

Cullin-RING ubiquitin ligases (CRLs) catalyze the ubiquitylation of substrates many of which are degraded by the 26S proteasome. Their modular architecture enables recognition of numerous substrates via exchangeable substrate receptors that competitively bind to a cullin scaffold with high affinity. Due to the plasticity of these interactions there is ongoing uncertainty how cells maintain a flexible CRL repertoire in view of changing substrate loads. Based on a series of in *vivo* and *in vitro* studies, different groups proposed that the exchange of substrate receptors is mediated by a protein exchange factor named cullin-associated and neddylation-dissociated 1 (Cand1). Here, we have performed quantitative mathematical modeling to support this hypothesis. To this end we first show that the exchange activity of Cand1 necessarily leads to a trade-off between high ligase activity and fast receptor exchange. Supported by previous *in vivo* studies we argue that this trade-off yields an optimal Cand1 concentration where the time scale for substrate degradation becomes minimal. In a second step we show through simulations that (i) substrates bias the CRL repertoire leading to preferential assembly of ligases for which substrates are available and (ii) differences in binding affinities create a temporal hierarchy for the degradation of substrates. Together, our results provide general constraints for the operating regimes of molecular exchange systems and suggest that Cand1 endows the CRL network with the properties of an “on demand” system allowing cells to dynamically adjust their CRL repertoire to fluctuating substrate abundances.

**Author summary:** Cullin-RING ubiquitin ligases (CRLs) are multisubunit protein complexes where exchangeable substrate receptors (SRs) assemble on a cullin scaffold to mediate ubiquitylation and subsequent degradation of a large variety of substrates. In humans there are hundreds of different CRLs having potentially thousands of substrates. Due to the high affinity of cullin-SR interactions, it has long been a mystery how cells would maintain flexibility to sample the entire SR repertoire in order to match fluctuating substrate loads. Recent experiments indicate that the exchange of different SRs is mediated by a novel protein exchange factor (Cand1). However, the proposed biochemical function of Cand1 as a promoter of CRL activity remained difficult to reconcile with previous reports of Cand1 acting as an inhibitor of CRL activity *in vitro*. Here we show that these two findings are not contradictory, but that the exchange activity of Cand1 necessarily leads to a trade-off between high ligase activity and fast receptor exchange which leads us to predict an optimal Cand1 concentration and a temporal hierarchy for substrate degradation. Our results support the view that Cand1 endows the CRL network with the flexibility of an “on demand” system where relative CRL abundances are dictated by substrate availability.

## Introduction

Cullin-RING ubiquitin ligases (CRLs) are modular protein assemblies that target cellular substrates for ubiquitylation which may alter the substrate’s activity or lead to its degradation by the 26S proteasome [1,2]. As such CRLs have been implicated in the regulation of numerous cellular processes which makes them attractive targets for the development of anti-cancer drugs [3,4]. The class of SCF (Skp1-Cul1-F-box) ubiquitin ligases represents the defining member of the CRL family [5,6]. SCF ligases consist of a cullin1 (Cul1) scaffold (Fig. 1A) with the RING finger protein Rbx1 (RING-box protein 1) bound to its C-terminal domain [7]. The latter acts as a binding site for an associated ubiquitin-conjugating enzyme (E2). Substrates to be ubiquitylated are recognized by dedicated F-box containing substrate receptors which bind to the N-terminal region of Cul1 via the Skp1 adapter protein. There are potentially 69 SCF complexes in humans. Since their total concentration exceeds that of the cullin scaffold [8] access of free SRs to Cul1 is under competition. Also, Cul1-SR binding appears to be extraordinarily tight [9] making spontaneous dissociation of preformed SCF complexes extremely unlikely and raising the question how access of different SRs to Cul1 is regulated in cells.

**Fig 1.**
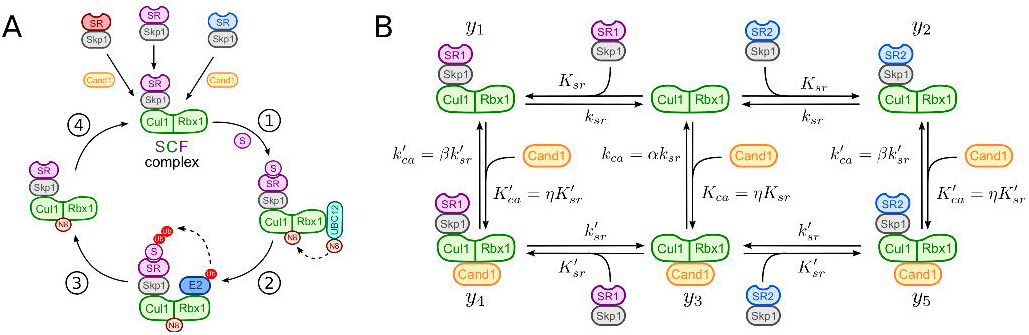
SCF-mediated substrate degradation and Cand1 cycle. **A**: Scheme of SCF-mediated substrate degradation: (1) Substrate (S) binding to substrate receptors (Skp1/SR) and UBC12-mediated neddylation (N8) of Cul1, (2) E2 recruitment, ubiquitin (Ub) transfer by E2 to the substrate and Ub chain elongation, (3) substrate degradation by the 26S proteasome and (4) deneddylation of Cul1 by the COP9 signalosome. Relative sizes of protein subunits are not to scale. **B**: Model of the Cand1-mediated exchange cycle for two substrate receptors (Skp1/SR1 and Skp1/SR2). *K_sr_*, *K_ca_*,
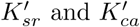
denote dissociation constants whereas *K_sr_*, *K_ca_*,
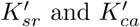
are dissociation rate constants (cf. Table 1). The parameter *η*, defined in Eq. (2), measures the preference of Cand1 and SR for binding to Cul1. Similarly, *α* and *β* account for relative differences in the dissociation rate constants for the binary complexes (*α*) and the ternary complex (*β*).

**Table 1.** Default parameter values.

| protein concentrations [nM] | dissociation constants [nM] | dissociation rate constants [$s^{-1}$] | scale factors (dimensionless) |
| --- | --- | --- | --- |
| $Cul1_T = 300$ <sup>(a)</sup> | $K_{sr} = 2.25 \cdot 10^{-4}$ <sup>(c)</sup> | $k_{sr} = 9 \cdot 10^{-7}$ <sup>(c)</sup> | $\eta = K_{ca}/K_{sr} = 0.077$ |
| $SR_T$ <sup>(b)</sup> = 660 <sup>(a)</sup> | $K'_{sr} = 650$ <sup>(c)</sup> | $k'_{sr} = 1.3$ <sup>(c)</sup> | $\alpha = k_{ca}/k_{sr} = 11.11$ |
| $Cand1_T = 390$ <sup>(a)</sup> | $K_{ca} = 1.73 \cdot 10^{-5}$ <sup>(d)</sup> | $k'_{ca} = 1 \cdot 10^{-5}$ <sup>(c)</sup> | $\beta = k'_{ca}/k'_{sr} = 0.031$ |
| | $K'_{ca} = 50$ <sup>(c)</sup> | $k'_{ca} = 0.04$ <sup>(e)</sup> | |

<sup>(a)</sup> Ref. [8], <sup>(b)</sup> $SR_T = SR1_T + SR2_T$ , <sup>(c)</sup> Ref. [9], <sup>(d)</sup> computed from Eq. (1), <sup>(e)</sup> cf. *Materials and Methods*.

SCF ligases are activated through covalent attachment of the ubiquitin-like protein Nedd8 to Cul1 which increases the binding affinity of Rbx1 for the E2 enzyme and promotes substrate ubiquitylation [10–12]. In the absence of substrate Nedd8 is removed from Cul1 by the COP9 signalosome (CSN) [8,13,14]. Interestingly, when Cul1 is not neddylated SRs can be exchanged by cullin-associated and neddylation-dissociated 1 (Cand1) which acts as an exchange factor for SRs [9,15,16]. Binding of Cand1 to a SCF complex dramatically increases the SR dissociation rate constant (10^6^-fold) similarly to nucleotide exchange factors when catalyzing the exchange of GDP for GTP in small GTPases [17], i.e. through formation of a short-lived ternary complex (Fig. 1B). Based on genetic evidence it has been argued that the exchange activity of Cand1 is required for efficient substrate degradation *in vivo* [18–21] and that Cand1 may potentially bias the assembly of SCF complexes towards F-box proteins for which substrates are available [9,22]. However, when analyzed *in vitro* Cand1 has been found to act as an inhibitor of SCF ligase activity [10,11,18,23–25]. In the present study we wish to show that these two findings are not contradictory, but that the exchange activity of Cand1 necessarily generates a trade-off between high SCF occupancy and fast SR exchange. As a result of this trade-off there exists an optimal Cand1 concentration where the time scale for substrate degradation becomes minimal. In a second step, we analyze the Cand1-mediated exchange of SRs in the presence of substrates which suggests a crucial role for Cand1 in shaping the cellular CRL repertoire.

## Models

### Model for the Cand1 exchange cycle

In our model (cf. Fig. 1B) we consider two species of substrate receptors (SRs) which competitively bind to the Cul1 scaffold via the adapter protein Skp1 [26]. Here, we do not explicitly model the assembly of Skp1 and SRs, but consider Skp1/SR dimers as preformed stable entities denoted for convenience by SR1 and SR2. Consistent with experiments we assume that binding of Cand1 to Cul1 lowers the binding affinity for SRs by 6 orders of magnitude [9], i.e.
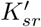
/*K*_sr_ ~ 10^6^, where *K_sr_* and
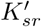
denote the dissociation constants of SRs from the binary Cul1.SR and the ternary Cul1.Cand1.SR complexes, respectively. Here and in the following we employ the “.” notation to denote non-covalent protein-protein interactions. Similarly, binding of SRs to Cul1 lowers the affinity for Cand1 by the same amount (i.e.
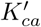
/*K_ca_* ~ 10^6^) resulting in the thermodynamic cycles depicted in Fig. 1B. Since the free energy change for the formation of the ternary Cul1.Cand1.SR complexes must not depend on the order in which they are formed the dissociation constants in each cycle have to satisfy the detailed balance relation

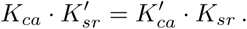

Here, *K_ca_* and
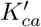
denote the dissociation constants of Cand1 from the binary Cul1.Cand1 complex and the ternary Cul1.Cand1.SR complexes, respectively. To satisfy the detailed balance condition (1) we introduce the relative binding affinity

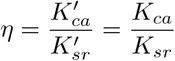

which measures the preference of Cand1 and SR for binding to Cul1, i.e. *η* < 1 means that Cand1 has a higher binding affinity for Cul1 whereas *η* > 1 means that SR has a higher binding affinity for Cul1.

To translate the reaction steps depicted in Fig. 1B into a mathematical model we employ mass-action kinetics and assume that the total protein concentrations (Cul1, Cand1, SR1 and SR2) remain constant on the time scale of interest. The dynamics of the system is described by 5 ordinary differential equations

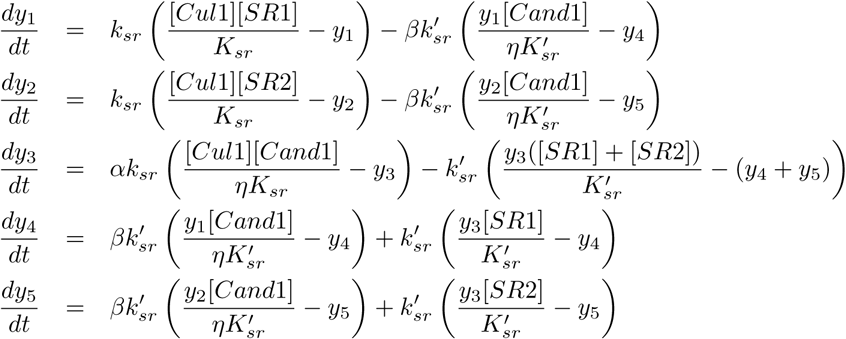

where *y*_1_,…, *y*_5_ denote binary and ternary protein complexes as indicated in Fig. 1B. The remaining variables in Eq. (3) can be expressed in terms of the *y_i_* using the mass conservation relations

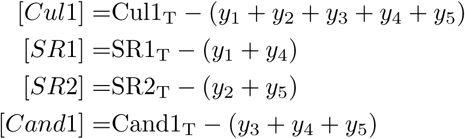

where Cul1_T_, SR1_T_, SR2_T_ and Cand1_T_ denote the total concentrations of Cul1, substrate receptors and Cand1, respectively.

To parametrize our model we employ representative rate constants as they were measured for the SCF^Fbxw7^ ligase (cf. Table 1). The only rate constant that remained undetermined in that study has been estimated from transient data in Ref. [9] (see *Materials and Methods*). Since all F-box containing SRs bind Cul1 via the Skp1 adapter protein we assume that both Skp1/SR dimers in Fig. 1B exhibit similar binding parameters as Skp1/Fbxw7. This idealized scenario is sufficient for our purpose as it allows studying competition effects between different SRs (through their relative abundances) while keeping the analysis of the system tractable. In Fig. 4 (below) we will relax this assumption and study the impact of different SR binding affinities on the time scale of substrate degradation.

## Results

### Cand1 reduces SCF ligase activity

We were first interested in understanding how the presence of Cand1 would affect the steady state occupancies for the different SCF complexes (i.e. Cul1.SR1 and Cul1.SR2). To this end, we assume that Cul1 is saturated with SRs, i.e. we consider the physiologically relevant regime SR_T_ = SR1_T_ + SR2_T_ > Cul1_T_. From the parameter values listed in Table 1 we see that *ηK_sr_* ≪ Cul1_T_. Under this condition we have derived approximate expressions for the steady state concentration of Cul1.SR1 (and the other complexes) in the limit of low and high concentrations of Cand1 (see Supporting Information S1 Text for details). In the first case (Cand1_T_ ≪ Cul1_T_) the SCF concentration decreases linearly with Cand1_T_ according to

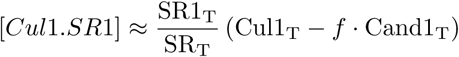

where the slope *f* is given by

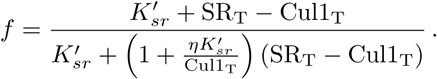

In contrast, for large Cand1 concentrations (Cand1_T_ ≫ CUl1_T_) the concentration of Cul1.SR1 decreases as a power law (~ 1/Cand1_T_) according to

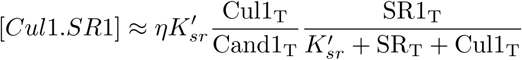

By symmetry, there exist similar expressions for [*Cul*1.*SR*2] with SR1_T_ being replaced by SR2_T_. Note that the slope parameter defined in Eq. (6) is limited to the range 0 < *f* < 1 where the lower (upper) bound is approached if
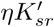
≫ Cul1_T_
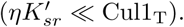
For the parameters listed in Table 1 *f* has the value 0.94 which is close to the upper bound. From Eqs. (5) and (7) we see that the SCF concentration decreases as a function of Cand1_T_ which is consistent with previous observations according to which Cand1 acts as an inhibitor of SCF ligase activity [18,23,24].

To analyze the behavior of the SCF occupancy near the transition point (where Cand1_T_ = Cul1_T_) we have plotted the steady state concentration of Cul1.SR1 for different values of the relative binding affinity *η* (Fig. 2A). We find that when *η* = 1 or larger the SCF concentration changes gradually near the transition point. However, when Cand1 exhibits a strong preference for binding to Cul1 (*η* ≪ 1) the SCF response curve develops a sharp threshold near the transition point (black line, Fig. 2A). Since the natural system seems to operate in the regime *η* ≪ 1 and Cand1_T_ > Cul1_T_ (cf. Table 1) one might expect that the concentration of SCF complexes (Cul1.SR1 and Cul1.SR2) is low under steady state conditions. However, this line of reasoning could be affected by two factors: First, the effective *in vivo* Cand1 concentration (available for binding to Cul1) could be lower than that of Cul1 because Cand1 also binds to other cullins of the CRL family [8]. Second, in the presence of substrates the concentration of particular SCF complexes could be increased due to dynamic remodeling of the SCF repertoire [7,9,22].

**Fig 2.**
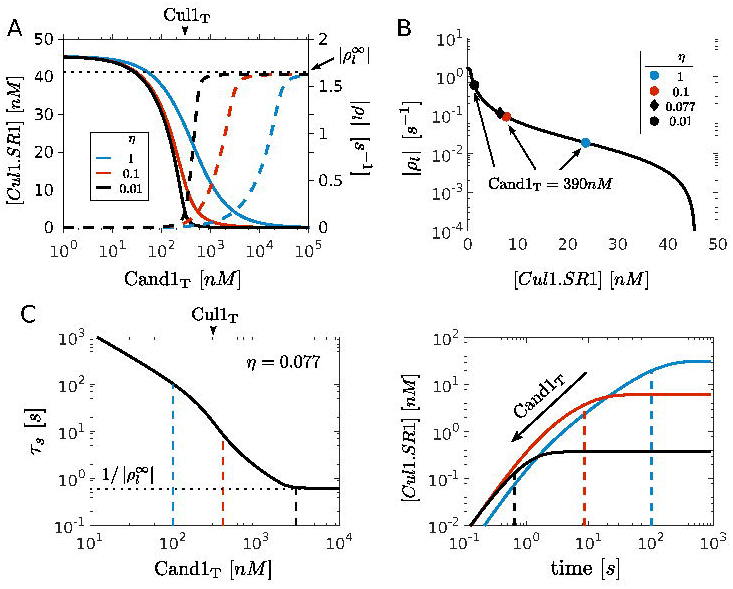
Trade-off between high SCF occupancy and fast SR exchange rate. **A**: Left axis shows SCF activity as measured by the steady state concentration of Cul1.SR1. Right axis shows the exchange rate as measured by the leading eigenvalue of the Jacobian matrix (*ρl*). As the total Cand1 concentration increases the SCF activity (solid lines) decreases while the exchange rate (dashed lines) concomitantly increases. As the relative binding affinity *η* (Eq. 2) decreases both the SCF response curve as well as the curve characterizing the exchange rate develop a sharp threshold near Cand1_T_ = Cul1_T_ (marked by arrow head). The horizontal dotted line indicates the maximal exchange rate (cf. Eq. 9). **B**: Exchange rate (|*ρl*|) vs. SCF occupancy ([*Cul*1.*SR*1]) drawn from the curves in panel A. Note that the curves are overlapping which suggests that the trade-off between high SCF occupancy and fast SR exchange rate is the same for all *η* ≤ 1. We have also indicated the positions along the curve where, depending on the value of *η*, the concentration of Cand1 equals that observed in cells (cf. Table 1). **C**: Left panel shows the time scale for the exchange of substrate receptors (*τ_S_* = 1/ |*ρ_l_*|) as a function of the total Cand1 concentration for the parameters listed in Table 1. Right panel shows the corresponding time course for the assembly of Cul1.SR1 after adding 100nM SR1 to a steady-state mixture containing 300nM Cul1, 560nM SR2 and Cand1 as indicated by the dashed lines in the left panel. The dashed lines in the right panel indicate the value of *τ_S_* obtained from the intersection of the dashed lines with the black solid line in the left panel.

### Trade-off between high SCF activity and fast SR exchange

Next we analyzed how Cand1 affects the time scale for the exchange of different SRs. If Cand1 is a SR exchange factor, as experiments suggest [9,15,16], the exchange rate should increase with increasing Cand1 concentration. To quantify the exchange rate we computed the leading eigenvalue of the Jacobian matrix (Fig. 2A and S1 Text) which determines the time scale for reaching a new steady state after applying a perturbation. Note that the SR exchange rate (as measured by |*ρl*|) dramatically increases when the Cand1 concentration is increased beyond that of Cul1 (the increase being more dramatic as *η* gets smaller). However, when Cand1_T_ is increased the SCF concentration ([*Cul*1.*SR*1]) concomitantly drops resulting in a trade-off between high SCF occupancy at low Cand1 concentration and fast SR exchange at high Cand1 concentration. This trade-off can be better visualized by plotting the SCF exchange rate against SCF occupancy (Fig. 2B). Interestingly, this yields the same curve independently of the value of *η*. However, depending on the value of *η* the Cand1 WT concentration of 390nM is reached at different positions along the trade-off curve (indicated by symbols). For example, in the WT system (diamond symbol) the SCF concentration is 6.4nM and the exchange rate is 0.11*s*^−1^.

To illustrate the impact of Cand1 on the time scale of SR exchange we assume that at *t* = 0 a fixed amount of SR1 is added to a steady state mixture of Cul1, Cand1 and SR2 with SR2_T_ > Cul1_T_ so that the cullin scaffold is already saturated with SRs prior to addition of SR1. After SR1 is added, a certain fraction of it gets exchanged for SR2 on Cul1. The time scale for the assembly of Cul1.SR1 ranges from a few minutes when Cand1_T_ ≪ Cul1_T_ to a few seconds when Cand1_T_ ≫ Cul1_T_ (cf. Fig. 2C).

To understand the constraints under which Cand1 mediates the exchange of SRs we consider again the two limiting regimes: Cand1_T_ ≪ Cul1_T_ and Cand1_T_ ≫ Cul1_T_. In the first case the leading eigenvalue can be approximated by (cf. S1 Text)

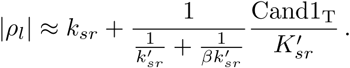

Consistent with expectation: As Cand1_T_ → 0, the SR exchange rate approaches the (spontaneous) dissociation rate constant of a Cul1.SR complex which is in the order of 10^−6^*s*^−1^ (cf. Table 1). In the presence of Cand1 the first term in Eq. (8) can be neglected showing that at low Cand1 concentrations the SR exchange rate is determined by the total rate with which the ternary complex dissociates into either of the two binary complexes, i.e. both branches (Cul1.Cand.SR → Cul1.SR and Cul1.Cand.SR → Cul1.Cand) contribute to the total dissociation rate. In contrast, if Cand1_T_ ≫ Cul1_T_ the SR exchange rate approaches a limiting value that is independent of *η* and Cand1_T_ (cf. Fig. 2A)

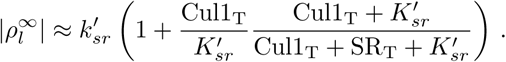

Since this expression depends
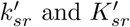
(but not on
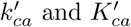)
it is the dissociation rate of the ternary complex towards Cul1.Cand1 which ultimately limits the rate with which new SRs can gain access to Cul1.

### Optimal Cand1 concentration

Due to the opposing effects of Cand1 on the SCF levels and the SR exchange rate we next asked how Cand1 would affect the substrate degradation rate. To this end, we extended the model depicted in Fig. 1B by assuming that substrate reversibly binds to Cul1.SR1 and that the substrate in the Cul1.SR1.S1 complex can be degraded by the proteasome (Fig. 3A). Here, we do not attempt to model the substrate degradation process in detail, instead we have lumped the relevant steps of the CRL cycle depicted in Fig. 1A (neddylation, ubiquitylation, deneddylation) into a single first order rate constant (*k_deg_*). To mimic the effect of neddylation we have assumed that once substrate is bound to its cognate SR the corresponding ligase complex becomes inaccessible for Cand1 so that SR exchange is suppressed [10,12]. Since Cand1 cannot bind to a neddylated SCF complex this assumption is consistent with the fact that substrate binding inhibits deneddylation and, therefore, favors the neddylated state [27,28]. To study the effects of Cand1 on the substrate degradation rate we performed numerical simulations where the total amount of SRs has been partitioned into 30nM SR1 and 630nM SR2. Then, a ten-fold excess of substrate for SR1 (SR1_T_ = 300*nM*) was added to a steady state mixture of Cul1, SR1, SR2 and different amounts of Cand1. Interestingly, the time scale for substrate degradation (measured by the time *t*_1/2_ it takes to degrade half of the total substrate) exhibited a non-monotonous behavior as a function of Cand1_T_ (Fig. 3B) changing from 48min (Cand1_T_ = 0*nM*) over 28min (Cand1_T_ = 100*nM*) to 95min (Cand1_T_ = 1000*nM*). Hence, there exists a minimum of *t*_1/2_ at intermediate Cand1 concentrations (Fig. 3C).

**Fig 3.**
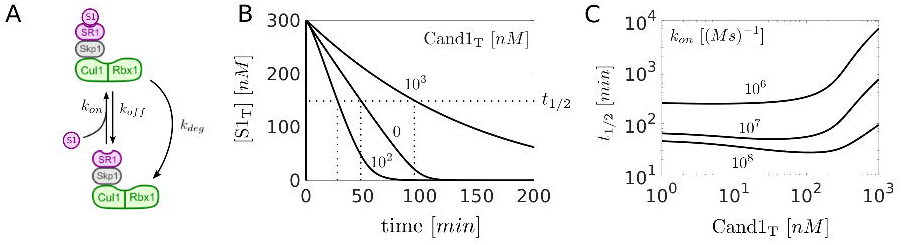
Optimal Cand1 concentration. **A**: Extension of the model depicted in Fig. 1B. Substrate (S1) reversibly binds to Cul1.SR1. The substrate in the resulting Cul1.SR1.S1 complex is degraded with effective rate constant *k_deg_*. **B:** Time courses showing the degradation of total substrate S_T_ = [*S*1] +[*Cul*1.*SR*1.*S*1]. At *t* = 0 substrate (300nM) was added to a steady state mixture containing SR1_T_ = 30*nM*, SR2_T_ = 630*nM* and Cand1_T_ as indicated. Note that *t*_1/2_ changes non-monotonically as a function of Cand1_T_. **C**: Half-life for substrate degradation (*t*_1/2_) as a function of Cand1_T_ for decreasing binding affinity. As the association rate constant *k_on_* for substrate binding decreases *t*_1/2_ increases and the dependence of *t*_1/2_ on Cand1_T_ becomes monotonous. Parameters for substrate binding: *k_on_* = 10^8^ *M*^−1^ *s*^−1^, *k_off_* = 1*s*^−1^, *k_deg_* = 0.004*s*^−1^. Parameters other than those mentioned are listed in Table 1.

The exchange of SRs by Cand1 takes time. So, we reasoned that Cand1 would lose its ability to accelerate substrate degradation if the binding affinity of the substrate for its cognate SR became too low. This is, indeed, what we observed (Fig. 3C): As the binding affinity decreases (*k_on_* decreases) the minimum vanishes and *t*_1/2_ increases monotonously with Cand1_T_ suggesting that Cand1 loses its ability to speed up substrate degradation for low-affinity substrates. Moreover, if Cand1_T_ becomes larger than Cul1_T_ = 300*nM* the *t*_1/2_ substantially rises independently of *k_on_* indicating that this regime might be unfavorable for efficient substrate degradation.

### Dynamic readjustment of the SCF repertoire

To increase the potential pool size of SCF ligases that can be engaged in the ubiquitylation of a cognate substrate unused SCF complexes should first be disassembled making the freed Cul1 available for the assembly of SCFs pertaining to the cognate substrate. This process is illustrated in Fig. 4: After addition of substrate the initial drop in [*Cul*1.*SR*1] is compensated by an increase in [*Cul*1.*SR*1.*S*1] (Fig. 4A). Later on, between 1-100min, the concentration of Cul1.SR1.S1 further rises due to disassembly of Cul1.SR2 and redistribution into Cul1.SR1 and Cul1.SR1.S1. The sum of the concentrations of these “engaged” SCF ligases ([Cul1.SR1] + [Cul1.SR1.S1]) increases 2.5-fold from its steady state value before it decreases back to pre-stimulus level after the substrate has been degraded (Fig. 4B, solid violet line). The remaining curves indicate the contribution from each of the other complexes (resulting from disassembly of Cul1.SR2, Cul1.SR1.Cand1, Cul1.SR2.Cand1 and Cul1.Cand1) assuming that redistribution of Cul1 occurred from only one of these complexes. Hence, at low Cand1 concentrations the majority of the redistributed Cul1 comes from Cul1.SR2 (Fig. 4B, blue line, long dashes) whereas at high Cand1 concentrations (Fig. 4C) the main contribution comes from Cul1.Cand1 (orange line, long dashes) and the ternary complexes (short dashes).

**Fig 4.**
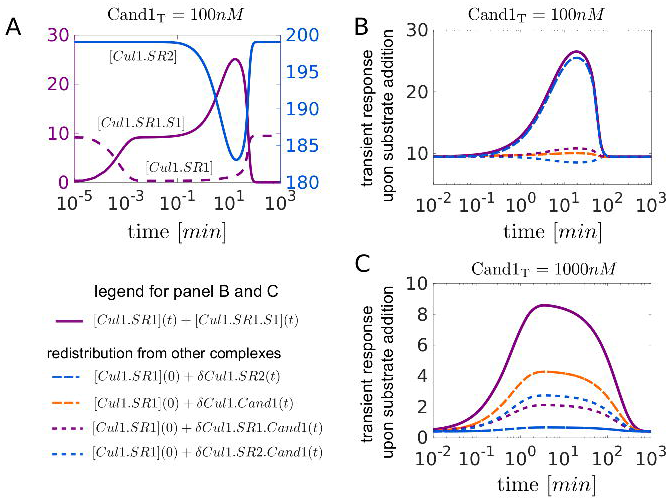
Dynamic readjustment of the SCF repertoire. **A**: Transient response of the SCF complexes Cul1.SR1, Cul2.SR2 and Cul1.SR1.S1 upon substrate addition (300nM at *t* = 0) to a steady state mixture containing SR1_T_ = 30*nM*, SR2_T_ = 630*nM* and Cand1_T_ = 100*nM*. Between 1-100min the drop in [Cul1.SR2] is accompanied by a peak in [Cul1.SR1.S1] indicating that Cul1 is redistributed from Cul1.SR2 into Cul1.SR1 and Cul1.SR1.S1. **B, C**: Redistribution of Cul1 from Cul1.SR2, Cul1.SR1(2).Cand1 and Cul1.Cand1 into Cul1.SR1 and Cul1.SR1.S1 for Cand1_T_ = 100*nM* (B) and Cand1_T_ = 1000*nM* (C). In both panels the solid violet line shows the transient increase of the concentration of “engaged” ligases ([Cul1.SR1]+[Cul1.SR1.S1]) upon substrate addition as described in (A). The remaining curves indicate the contribution to the transient response by any of the other complexes. For example, *δCul*1.*SR*2(*t*) = [*Cul*1.*SR*2](0) – [*Cul*1.*SR*2](*t*) denotes the amount of Cul1 that is redistributed into Cul1.SR1 and Cul1.SR1.S1 upon disassembly of Cul1.SR2. Note that in each of the panels B and C the solid violet curve is the sum of the other curves. Parameters for substrate binding: *k_on_* = 10^8^*M*^−1^*s*^−1^, *k_off_* = 1*s*^−1^, *k_deg_* = 0.004*s*^−1^. Parameters other than those mentioned are listed in Table 1.

### Temporal hierarchy of substrate degradation

Another interesting question is whether there exists a temporal order in which SR substrates are degraded by the 26S proteasome. To analyze such a scenario we extended the model depicted in Fig. 1B and considered 3 types of SRs: two for which substrates are available (SR1 and SR2) and one representing the remaining SR pool (SR3). It is assumed that downstream processing by the proteasome is the same for both substrates (*k_deg_* = 0.004*s*^−1^), but that there might be differences in the binding affinity of substrate to their cognate SR (Fig. 5A,D), differences in the binding affinity of SRs to Cul1 (Fig. 5B, E) or differences in SR abundances (Fig. 5C, F). Our simulations suggest that differences in either of these parameters can induce a temporal order in the degradation of substrates such that high-affinity substrates, substrates with high-affinity SRs and substrates of highly abundant SRs are degraded first (as indicated by a lower *t*_1/2_). In all cases substrate degradation is accompanied by a redistribution of Cul1 from the pool of unused SCFs (SCF3) into the pool of engaged SCFs (SCF1 and SCF2) supporting the view that in the presence of substrates the exchange activity of Cand1 leads to the preferential assembly of SCFs for which substrates are available [9,22].

**Fig 5.**
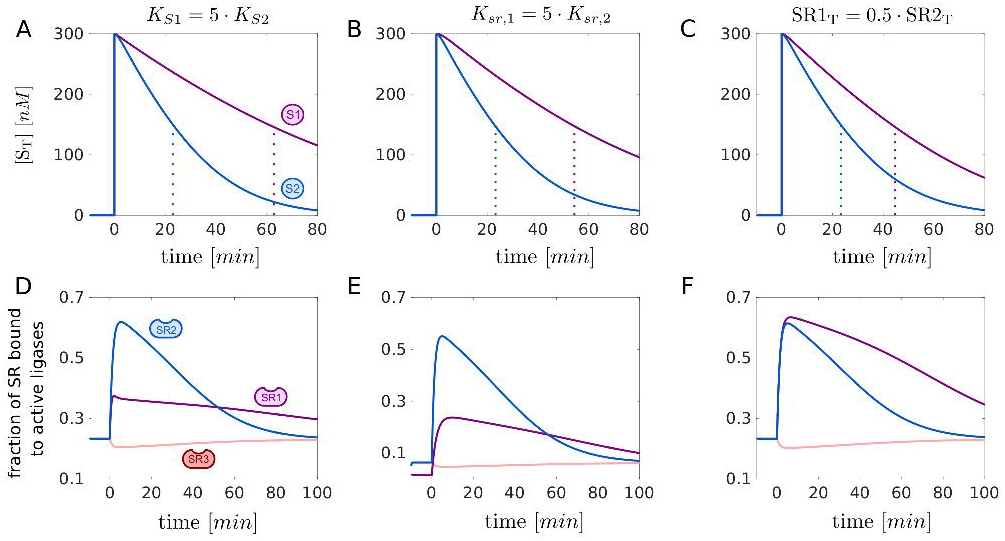
Temporal hierarchy of substrate degradation. **A**, **B**, **C**: Transient response upon substrate addition. At *t* = 0 two substrates, S1 and S2 (each 300nM), are added to a steady state mixture containing Cul1, Cand1 and SR1-SR3. The resulting decline of the total amount of substrates is displayed together with the *t*_1/2_ (dotted lines). Substrates with a higher SR affinity (A), substrates for SRs with a higher affinity for Cul1 (B) and substrates for more abundant SRs (C) are preferentially degraded. **D, E, F**: Assembly and disassembly of SCF ligases upon substrate addition. Depicted are changes in the fraction of SRs that are bound in a SCF complex. The blue and violet curves correspond to ([*Cul*1.*SR*1] + [*Cul*1.*SR*1.*S*1])/SR1_T_ and ([*Cul*1.*SR*2] + [*Cul*1.*SR*2.*S*2])/SR2_T_, respectively, whereas the light red curve denotes [*Cul*1.*SR*3]/SR3_T_. In each case Cul1 is redistributed from Cul1.SR3 into Cul1.SR1(.S1) and Cul1.SR2(.S2). In (A-F) if not indicated otherwise reference parameters are: *K*_*S*1_ = *K*_*S*2_ = 10*nM* (*k_off_* = 1*s*^−1^), *K*_*sr*,1_ = *K*_*sr*,2_ = *K*_*sr*,3_ = 0.225*pM*, SR1_T_ = SR2_T_ = 60*nM*. To preserve detailed balance
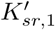
has been increased by a factor of 5 in (B) and (E). SR3_T_ = 660*nM* – (SR1_T_ + SR2_T_), Cand1_T_ = 400*nM*, *k_deg_* = 0.004*s*^−1^. Parameters other than those mentioned are listed in Table 1.

### Is Cand1 necessary for fast SR exchange?

One of the puzzling properties of SCF ligases (and perhaps other CRLs) is the extremely high affinity of the Cul1-SR interaction which lies in the picomolar range [9]. One reason might be to prevent “leakage” so that SR exchange is exclusively mediated by Cand1. Indeed, experiments with *cand1* deletion cell lines have shown that most F-box proteins rely on the exchange activity of Cand1 for efficient substrate degradation [15,16,21]. Alternatively, one could envision a system with substantially weaker Cul1-SR interaction. In such a hypothetical system newly synthesized F-box proteins could always gain access to Cul1 making an exchange factor dispensable.

To compare these two architectures we rescale the dissociation rate constant *k_sr_* by a factor *γ* > 1 which lowers the binding affinity between Cul1 and SR (Fig. 6A). To satisfy the detailed balance condition in Eq. (1) we multiply *k_ca_* by the same factor so that the dissociation constants *K_sr_* and *K_ca_* increase with *γ* while their ratio remains constant. In this setting the case *γ* =1 and Cand1_T_ = 390*nM* corresponds to the wildtype system whereas the case *γ* > 1 and Cand1_T_ = 0*nM* represents the alternative system design. To make a fair comparison we chose *γ* such that the steady state level of Cul1.SR1 prior to addition of substrate (S1) is the same for both cases. In addition, we assume that substrate can bind to both Cul1.SR1 and Cul1.Cand1.SR1. To mimic the effect of neddylation in this setting we allow substrate to be degraded only when it is bound to Cul1.SR1, but not when it is bound to Cul1.Cand1.SR1 (since Cand1 and Nedd8 cannot be simultaneously bound to Cul1). Interestingly, the half-life of substrate degradation depends not only on the presence or absence of Cand1, but also on the detailed mechanism of substrate binding: If substrate can only bind to SR when the latter is already bound to Cul1 or Cul1.Cand (sequential mechanism) the system without Cand1 exhibits faster substrate degradation (3.4-fold) compared to the system with Cand1 (Fig. 6B). In contrast, when substrate binding occurs in a random manner the situation is reversed as substrate degradation is now faster (4.1-fold) in the presence of Cand1 (Fig. 6C).

**Fig 6.**
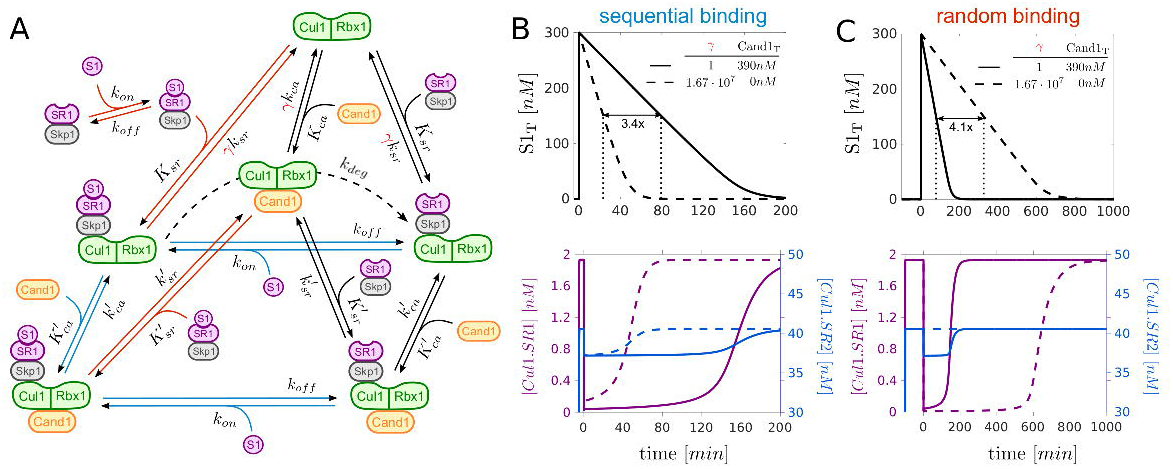
Alternative network architecture. **A**: Extension of the Cand1 cycle model for SR1 (black solid lines) to include substrate binding. Sequential mechanism: Substrate (S1) only binds to SR1 if the latter is already bound to Cul1or Cul1.Cand1 (blue lines). Random mechanism: S1 also binds to free SR1, Cul1.SR1 and Cul1.Cand1.SR1. In addition, SR1.S1 binds to Cul1 or Cul1.Cand1 (red lines). By increasing the factor *γ* (red color) the binding affinity between Cul1 and SR1 can be lowered while still satisfying the detailed balance condition in Eq. 1. For SR2 we use the same scheme as depicted in Fig. 1B (without substrate) with *k_sr_* and *k_ca_* multiplied by *γ*. *K_sr_*, *K*_ca_,
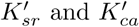
denote dissociation constants whereas *k_sr_*, *k*_ca_,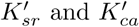
are dissociation rate constants (cf. Table 1). **B**: Comparison of the half-life of S1 (*t*_1/2_) for two network designs: one with Cand1_T_ > 0 and tight binding of SRs to Cul1 (*γ* = 1) and another one with Cand1_T_ = 0 and weak binding of SRs to Cul1 (*γ* ≫ 1). In the latter case *γ* is chosen such that the pre-stimulus steady state for Cul1.SR1 is the same in both cases (Note that dashed and solid lines in lower panels partially overlap). If substrate binds sequentially the system with Cand1_T_ = 0*nM* (B, dashed line) outperforms the system with Cand1_T_ = 390*nM* (B, solid line) as the *t*_1/2_ is 3.4-fold larger in the presence of Cand1. In both cases Cul1 is redistributed from Cul1.SR2 to Cul1.SR1 and Cul1.SR1.S1 (B, lower panel). In contrast, when substrate binds in a random manner (cf. panel A) its degradation is substantially delayed (4.1-fold) in the absence of Cand1 (C) and redistribution of Cul1 only occurs in the presence of Cand1 (C, lower panel). Total substrate is defined as S1_T_ = [*S*1] + [*SR*1.*S*1] + [*Cul*1.*SR*1.*S* 1] + [*Cu1*1.*Cand*1.*SR*1.*S*1]. Parameters: A*t t* = 0 substrate S1 (300*nM*) was added to a steady state mixture containing Cul1_T_ = 300*nM*, SR1_T_ = 30*nM* and SR2_T_ = 630*nM*. The values of Cand1_T_ and *γ* are indicated in the upper panels. *k_on_* = 10^7^*M*^−1^ *s*^−1^, *k_off_* = 0.01*s*^−1^, *k_deg_* = 0.004*s*^−1^. Parameters other than those mentioned are listed in Table 1.

There are two factors that might explain this behavior: First, for the system without Cand1 redistribution of Cul1 from Cul1.SR2 into Cul1.SR1 and Cul1.SR1.S1 only occurs if substrate binding occurs sequentially (Fig. 6B, C lower panels). Second, when binding occurs randomly in a system without Cand1 substrate may become “trapped” in SR1.S1 complexes which bind only weakly to Cul1. Since in such a system the Cul1-SR binding affinity (*γK_sr_* ≈ 37*nM*) is weaker than the assumed substrate affinity (1nM) binding to free SRs effectively reduces the substrate’s affinity for gaining access to Cul1 which causes a delay in its degradation.

## Discussion

Cullin-RING ubiquitin ligases (CRLs) are multisubunit protein complexes where exchangeable substrate receptors (SRs) assemble on a cullin scaffold to mediate ubiquitylation and subsequent degradation of a large variety of substrates. Motivated by the observation that the exchange of different SRs is catalyzed by an exchange factor (Cand1) [9,15,16] we were interested in the operating regimes and the inherent constraints that may exist in such exchange systems, and how they would affect the degradation of ubiquitylation substrates. Specifically, we wanted to understand how the CRL network can flexibly react to changingsubstrate loads despite the high-affinity of cullin-SR interactions.

Our results indicate that there exists a generic trade-off in the Cand1-mediated exchange of SRs which leads to an optimal Cand1 concentration where the time scale for substrate degradation becomes minimal (cf. Fig. 3C). This result can be rationalized as follows: In the absence of Cand1 only preassembled SCF complexes contribute to substrate degradation since free SRs cannot gain access to Cul1. As the Cand1 concentration increases the concentration of preassembled SCF complexes decreases since part of the Cul1 is sequestered by Cand1 into Cul1.Cand1 and ternary Cul1.Cand1.SR complexes, which are necessary to mediate the exchange of SRs. However, in the presence of Cand1 disassembly and reassembly of SCFs increases the effective pool size of SCF ligases for a particular substrate at the expense of unused SCF ligases which more than compensates the drop of preassembled SCFs and reduces the time scale for substrate degradation. If, on the other hand, the Cand1 concentration becomes substantially larger than that of Cul1 sequestration of Cul1 into Cul1.Cand1 and ternary complexes dominates. In this regime the drop of preassembled SCFs cannot be compensated anymore by the increased exchange activity of Cand1 resulting in an increased time scale for substrate degradation. Together, these results show that, by lowering the SCF occupancy, the exchange activity of Cand1 necessarily leads to an apparent reduction of SCF ligase activity which is consistent with previous reports of Cand1 acting as an inhibitor of SCF ligases [10,11,18,23–25]. Our results also support the view that in the presence of SCF substrates the exchange activity of Cand1 may bias the SCF repertoire leading to the preferential assembly of SCF ligases for which substrate is available (Fig. 4) [9,22]. As such Cand1 may endow the CRL network with the flexibility of an “on demand” system, thereby allowing cells to dynamically adjust their CRL repertoire to fluctuating substrate abundances.

Experimental evidence for an optimal Cand1 concentration *in vivo* comes from experiments by Lo and Hannink [21] who found (in two different cell lines) that both overexpression of Cand1 as well as siRNA-mediated knockdown of Cand1 leads to increased steady state levels of the transcription factor Nrf2. This is consistent with our finding that increasing and lowering the Cand1 concentration beyond and below the optimal level (where the half-life is minimal) leads to an increased half-life of substrates. Nrf2 is an ubiquitylation target of the Cul3-Keap1 ubiquitin ligase whose assembly has been shown to be controlled by Cand1 [29]. This suggests that our results may not only apply to SCF ligases, but also to other members of the CRL family. Based on the measured rate constants listed in Table 1 our model predicts an optimal Cand1 concentration in the range between 30nM – 120nM depending on the substrate’s binding affinity. When comparing this prediction with the cellular concentrations of Cand1 (390nM) and Cul1 (302nM) one has to take into account that Cand1 not only binds to Cul1, but also to cullins of other CRL family members (Cul2-Cul5) whose total concentration adds up to ≈ 1260nM [8]. Hence, the *in vivo* Cand1/CRL ratio of ~ 0.3 falls onto the upper boundary of the predicted range of optimal Cand1 concentrations indicating that in cells the exchange activity of Cand1 might be optimized for high-affinity substrates. In fact, our simulations show that Cand1 loses its ability to speed up substrate degradation when the substrate’s binding affinity becomes too low (Fig. 3C). In addition, substrate degradation is predicted to occur in a temporal order with high-affinity substrates being degraded first (Fig. 5). Similar effects are seen for high-affinity and highly abundant SRs suggesting that cells may exploit several mechanisms to fine-tune substrate degradation to needs.

From a mechanistic point of view the Cand1-mediated exchange of SRs exhibits some similarity to the exchange of GDP by GTP as mediated by guanosine nucleotide exchange factors (GEFs) [17]. However, while GEFs catalyze the exchange between only two substrates, Cand1 potentially mediates the exchange of hundreds of different SRs. When comparing the parameters of the Cand1 cycle with those of GDP/GTP exchange cycles one finds several systems that seem to operate in a similar regime. For example, in the Ran/RCC1 as well as in the EF-Tu/EF-Ts systems the concentration of the exchange factors, RCC1 and EF-Ts, is typically lower than that of the respective GDP/GTP-binding proteins [30,31]. Also, the binding affinities of GDP and the exchange factor with respect to EF-Tu or Ran are either comparable [32] or there exists a slight preference in favor of the exchange factor [30] suggesting that both systems operate in the regime *η* ≤ 1. Similar as for the Cand1 cycle this may indicate that the concentration of the respective exchange factor is optimized for the purpose of the system, e.g. fast nuclear export rate of proteins in the case of Ran/RCC1 and a high protein synthesis rate in the case of EF-Tu/EF-Ts. Indeed, theoretical studies have shown that GDP/GTP exchange systems potentially exhibit similar trade-offs as the ones reported here for the Cand1 cycle [33,34] although direct experimental evidence for an optimized concentration of the exchange factor seems to be lacking in those cases.

## Materials and Methods

The simulations depicted in Fig. 2 were done using MatCont [35]. The transient simulations involving substrate degradation depicted in Figs. 3-5 were done using the SimBiology Toolbox of MATLAB [36]. Derivations of the analytical formulas in Eqs. 5 - 9 can be found in S1 Text.

### Model extensions: Substrate binding and 3 substrate receptors

To generate the simulations depicted in Fig. 3 and Fig. 4 we have assumed that substrate (S1) reversibly binds to its cognate substrate receptor (Skp1/SR1) with forward and backward rate constants *k_on_* and *k_off_*. Not much seems to be known about the values of these parameters for particular substrates, so we set *k_on_* = 10^8^*M*^−1^*s*^−1^ (close to the diffusion limit) and *k_off_* = 1*s*^−1^ giving a binding affinity of *K_D_* = *k_off_*/*k_on_* = 10*nM*. Given the tight binding between Cul1 and SRs (*K_D_* ~ 1*pM*) it seems likely that typical binding affinities between substrates and their cognate receptors are even lower than 10nM. Substrate degradation has been modeled through a first order process of the form *Cul*1.*SR*1.*S* 1 → *Cul*1.*SR*1 with an effective first order rate constant of *k_deg_* = 0.004*s*^−1^ (corresponding to 0.24*min*^−1^).

To conduct the simulations shown in Fig. 5 we have considered two substrates (S1 and S2) each binding to its cognate SR with the same set of default values for *k_on_* = 10^8^*M*^−1^*s*^−1^, *k_off_* = 1*s*^−1^ and *k_deg_* = 0.004*s*^−1^. In addition, we have included another substrate receptor species (SR3) which collectively accounts for auxiliary receptors that compete for access to Cul1, but for which no substrate is available. To this end, we have added three reversible binding equilibria similar to those already depicted for SR1 and SR2 assuming for each of the reactions the same value for *kon* and *k_off_* as for SR1 and SR2. To generate the curves in Figs. 5A, D we have lowered the *k_on_* for S1 5-fold to *k_on_* = 2 ⋅ 10^7^*M*^−1^*s*^−1^ so that *K*_*D*,*S*1_ = 5 ⋅ *K*_*D*,*S*2_. To generate the curves in Figs. 5B, E we have increased *k*_*sr*,1_ for SR1 5-fold to *k*_*sr*,1_ = 4.5 ⋅ 10^−6^*s*^−1^ so that the Cul1-binding affinity of SR1 is 5-fold lower compared to that of SR2, i.e. *K*_*sr*,1_ = 5 ⋅ *K*_*sr*,2_. To preserve the detailed balance relation (Eq. 1) for the cycle involving SR1 we have also increased the value of
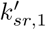
5-fold to
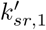
= 6.5*s*^−1^.

### Estimation of 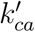

The rate constants listed in Table 1 were measured for the particular substrate receptor Fbxw7 using a FRET-based assay [9]. The dissociation constants *K_sr_* and
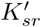
were directly computed from the reported values for *k_on_* and *k_off_*. For the dissociation constant
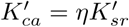
an upper limit of 50nM has been reported. Using this value as an estimate for
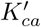
yields *η* ≈ 0.077. The remaining dissociation constant is then determined by the detailed balance relation (Eq. 1) which yields *K_ca_* ≈ 1.73 ⋅ 10^−5^*nM*. From the 4 dissociation rate constants listed in Table 1 only
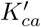
had not been measured. To estimate this parameter we repeat the experiment in Fig. 4B from Ref. [9] in a computer simulation (Fig. 7A). Here, 70nM of CFP-tagged Cul1 was first incubated with 70nM *β*-TrCP-Skp1 which sequestered essentially all the available Cul1. Hence, subsequent addition of TAMRA-labelled Fbxw7 (Fbxw7^TAMRA^–Skp1) did not evoke a change in donor fluorescence since Fbxw7-Skp1 could not gain access to Cul1. However, in the presence of 150nM Cand1 the fluorescence signal decayed over time due to Cand1-mediated exchange of *β*-TrCP for Fbxw7. An exponential fit to the time course of the signal yielded a first order rate constant of *k_obs_* ≈ 0.07*s*^−1^. In our model we have changed
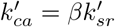
(the only free parameter) until the time scale for reaching the steady state coincided with that observed experimentally (Fig. 7B). As a result we obtained
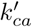
= 0.04*s*^−1^ or *β* = 0.031.

**Fig 7.**
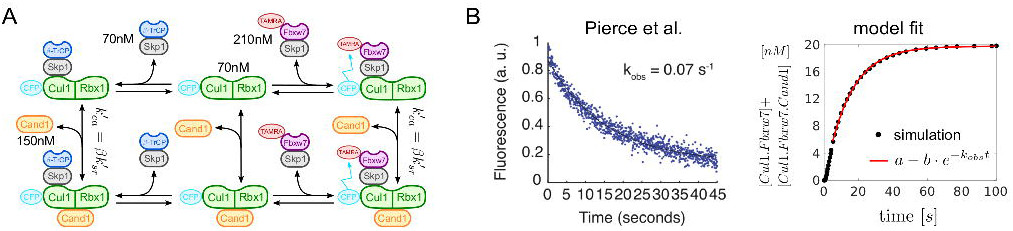
Estimation of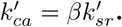. **A**: Scheme showing the reactions as used in the experimental setup of Pierce et al. [9]. States and reactions have the same meaning as in Fig. 1B. Note that Cul1 and Fbxw7 are labeled by fluorescent dyes. **B:** Left panel shows the change in donor fluorescence upon binding of Fbxw7^TAMRA^-Skp1 to ^CFP^Cul1-Rbx1. The picture has been taken from Fig. 4B of Pierce et al. [9]. The right panel shows the fit of the model simulations to a single exponential function. Data points for the first 5 seconds were discarded to obtain a better fit.

## Supporting information

**S1 Text Steady state and time scale analysis of the Cand1 cycle model**. In S1 Text we conduct a steady state/time scale analysis of Eqs. (3) and provide the derivations of Eqs. (5) - (9).

## Acknowledgments

We thank Michal Sharon and Wolfgang Dubiel for comments on the manuscript. RS is grateful for discussions with Ray Deshaies. This work was supported by grant GM105802 to DAW. Parts of this work were also funded by P30 grants CA030199 and GM085764. DAW is a scholar of the Foreign 1000 Talent Program funded by the Government of the People’s Republic of China. DF is deeply grateful for the support by the Max Planck Institute for Dynamics of Complex Technical Systems and for support by the Institute for Automation Engineering (IFAT) at the Otto-von-Guericke-University Magdeburg.

